# Functional Anabolic Network Analysis of Human-associated *Lactobacillus* Strains

**DOI:** 10.1101/746420

**Authors:** Thomas J. Moutinho, Benjamin C. Neubert, Matthew L. Jenior, Maureen A. Carey, Gregory L. Medlock, Glynis L. Kolling, Jason A. Papin

**Affiliations:** Department of Biomedical Engineering, University of Virginia, Charlottesville, Virginia, USA; Division of Infectious Disease and International Health, Department of Medicine, University of Virginia, Charlottesville, Virginia, USA

## Abstract

Members of the *Lactobacillus* genus are frequently utilized in the probiotic industry with many species conferring demonstrated health benefits; however, these effects are largely strain-dependent. We designed a method called PROTEAN (Probabilistic Reconstruction Of constituent Anabolic Networks) to computationally analyze the genomic annotations and predicted metabolic production capabilities of 144 strains across 16 species of *Lactobacillus* isolated from human intestinal, oral, and vaginal body sites. Using PROTEAN we conducted a genome-scale metabolic network comparison between strains, revealing that metabolic capabilities differ by isolation site. Notably, PROTEAN does not require a well-curated genome-scale metabolic network reconstruction to provide biological insights. We found that predicted metabolic capabilities of lactobacilli isolated from the vaginal microbiota cluster separately from intestinal and oral isolates, and we also uncovered an overlap in the predicted metabolic production capabilities of intestinal and oral isolates. Using machine learning, we determined the most informative metabolic products driving the difference between predicted metabolic capabilities of intestinal, oral, and vaginal isolates. Notably, intestinal and oral isolates were predicted to have a higher likelihood of producing D-alanine, D/L-serine, and L-proline, while the vaginal isolates were distinguished by a higher predicted likelihood of producing L-arginine, citrulline, and D/L-lactate. We found the distinguishing products to be consistent with published experimental literature. This study showcases a systematic technique, PROTEAN, for comparing the predicted functional metabolic output of microbes using genome-scale metabolic network analysis and computational modeling and provides unique insight into human-associated *Lactobacillus* biology.

**Importance:** The *Lactobacillus* genus has been shown to be important for human health. Lactobacilli have been isolated from human intestinal, oral, and vaginal sites. Members of the genus contribute significantly to the maintenance of vaginal health by providing colonization resistance to invading pathogens. A wide variety of clinical studies have indicated that *Lactobacillus*-based probiotics confer health benefits for several gut- and immune-associated diseases. Microbes interact with the human body in several ways, including the production of metabolites that influence physiology or other surrounding microbes. We have conducted a strain-level genome-scale metabolic network reconstruction analysis of human-associated *Lactobacillus* strains, revealing that predicted metabolic capabilities differ when comparing intestinal/oral isolate to vaginal isolates. The technique we present here allows for direct interpretation of discriminating features between the experimental groups.

## Introduction

*Lactobacillus* is a diverse genus of bacteria with many member strains associated with the human body. Lactobacilli are Gram-positive, lactic acid-producing bacteria typically with a low GC content (1,2). They are known for their production of lactic acid, being facultative anaerobes, and are capable of being metabolically active in a large variety of conditions (3). There is evidence that human-associated lactobacilli colonize mucosal surfaces of the intestinal tract (4), vagina (5–12), and oral cavity (13,14). While strains of *Lactobacillus* have been isolated from all three of these body sites, it remains unknown which are permanent members of the resident microbiota (autochthonous) opposed to transient members (allochthonous). Transient intestinal lactobacilli are either resident members of the oral microbiota or have been ingested, most commonly from unpasteurized fermented foods (4,15).

Lactobacilli have been used for a broad range of applications primarily associated with human intestinal probiotics and industrial production of useful metabolites. *Lactobacillus*-based probiotics have been shown to confer health benefits in clinical studies for a variety of conditions including prevention of antibiotic associated diarrhea (16), *Clostridium difficile-*associated diarrhea (17), constipation (18), irritable bowel syndrome (19), and eczema/atopic dermatitis (20). Probiotics are controversial, likely due to claims made by currently marketed probiotics that lack FDA approval for the treatment of specific diseases (21,22). The primary benefits associated with lactobacilli-based probiotics may be a function of their presence in the gut, production of metabolites, and modulation of the immune system (23,24). Metabolism plays a key role in all three of these general mechanisms; therefore, a better understanding of their metabolic capabilities will help to elucidate the mechanisms contributing to probiotic effects (25).

In recent years, there has been an explosion of genomic and metagenomic sequencing of human-associated microbiota, which provides a unique opportunity to apply genome-scale metabolic network reconstructions (GENREs) to enhance our current understanding of human-associated lactobacilli metabolism utilizing *in silico* techniques (25). Systems biology has the potential to advance design, selection, and delivery of *Lactobacillus*-based probiotics (26,27). GENREs are a powerful computational tool for mathematically modeling the metabolic processes within a cell at a systems-level, including all known metabolic reactions, metabolites, and metabolic genes in an organism (28). GENREs are created by referencing an annotated genome against biochemical databases, then integrating experimental data when available (29). There are several examples of *Lactobacillus*-specific comparative genomics studies (30–35); however, GENREs allow for a more functional perspective than genomics data alone because of the quantitative accounting for interactions between components in the network (25,36). Simulations with GENREs can accurately predict microbial growth yields and the metabolic pathways utilized for the production of metabolites during exponential growth of a microbe (37). A variety of analytical approaches can be applied to interrogate emergent properties of a GENRE. Flux Balance Analysis (FBA) and related methods have proven highly successful in the analysis of metabolic networks (38). FBA is a mathematical technique for analyzing the flow of metabolites through a GENRE; it can be used to identify a set of reaction fluxes that maximize growth in a specified media condition among other applications (28,39,40). Metabolic network reconstructions and FBA provide a mechanistic look into cellular metabolism and are increasingly used to study biochemical processes of single bacterial species as well as communities of organisms (41).

GENREs enable the computational prediction of metabolic capabilities of microbes, both catabolic and anabolic. Additionally, GENREs are capable of contextualizing large ‘omic datasets (i.e. genomics, transcriptomics, and metabolomics) with known biochemistry and biological network architectures for improved understanding of the experimental data (42). An important recent finding demonstrated that metabolomics data alone can be used to differentiate between bacterial cultures at the strain level (43). We developed a computational method using GENREs to predict the metabolic products that a strain is likely able to produce. We used predicted production capabilities to then differentiate between different human-associated *Lactobacillus* strains. Just as metabolomics data can be used to differentiate bacterial strains, predicted production capabilities can be used for the same comparisons. We assessed the metabolic potential across a broad set of *Lactobacillus* species, consisting of 144 strains, which have all been isolated from three human-related body sites: intestinal, oral, and vaginal. We found that intestinal and oral isolates have a great deal of overlap in their metabolic functionality, while vaginal isolates have more unique metabolic production capabilities. These analyses can facilitate additional experimental interrogation of this important genus of bacteria.

## Results and Discussion

### Annotated metabolic genes associated with known metabolic functions are sufficiently represented among human-associated lactobacilli

In this study we predict the metabolic production capabilities of 144 lactobacilli strains. We utilized the PATRIC Cross-Genus Protein Families (PGfams) (4) for an initial genomic analysis. PGfams are comparable clusters of proteins that likely have similar functions. These clusters are intended to be used for cross-genus comparison due to their slightly relaxed clustering criteria. However, PGfams allow for the comparison of the large number of strains analyzed in this study. Lactobacilli consist of a broad range of species and thus using the PGfams was appropriate for an initial genomic comparison in this study. We first filtered the PGfams to only include metabolic gene families associated with known metabolic functions (see Methods). The distribution of total metabolic PGfams associated with each genome ranges from 340 to 580 and has a median value of 515 (Figure 1A). Across these 144 strains we found that they share 116 core metabolic PGfams, spanning a variety of cellular functions including, but not limited to, carbohydrate, nucleotide, and amino acid metabolism (Table S1). The pan set of metabolic PGfams, which represents the total set of unique PGfams, expanded to over 1500 after considering all strains utilized within this study (Figure 1B). The *Lactobacillus* strains we studied consisted of 16 species and were isolated from intestinal, oral, and vaginal human body sites (Figure 1C).

**Figure 1:**
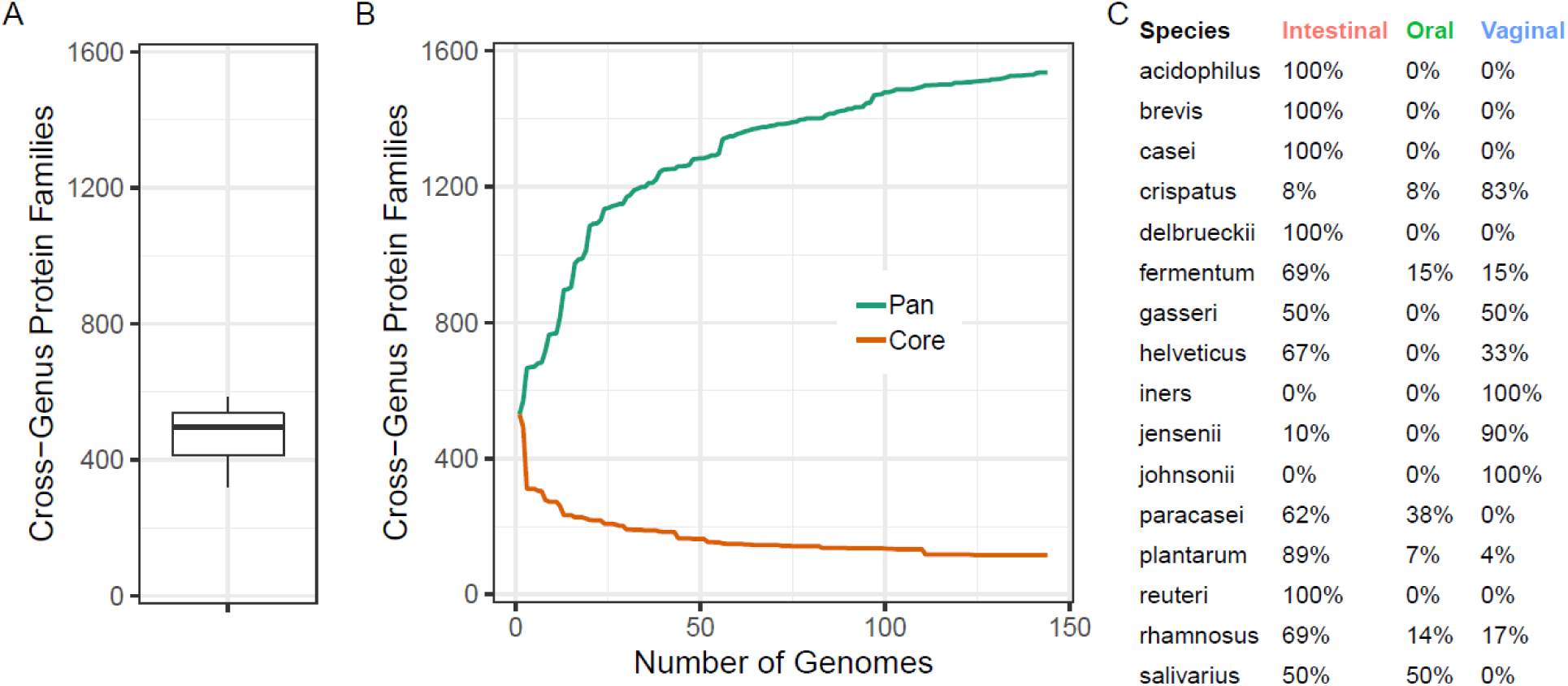
Known metabolic annotations are extensively sampled across the 16 *Lactobacillus* species included in this study. The genomic features used for this analysis are PATRIC Cross-Genera Protein families (PGfams), a standardized set of features across the PATRIC Database (4). (A) The number of metabolic PGfams for each genome are shown here, with the median value indicated by the middle line in the boxplot. (B) For the 144 strains from 16 species of *Lactobacillus*, we found that there are 116 protein families in the core set of metabolic PGfams, while the pan set of PGfams expands to over 1500 families. The nearly plateau shape of the curve for the pan set of PGfams curve indicates that this sampling represents a large portion of the genetic diversity among the 16 species included in the study. (C) This table shows the complete list of species used in this study and indicates the percentage of strains that were isolated from each human body site. Each strain in this study is a member from one of the 16 species and isolated from one of three human-associated body sites; intestinal, oral, or vaginal (Table S2).

### Probabilistic Reconstruction Of constituent Anabolic Networks (PROTEAN)

We developed PROTEAN to predict the metabolic production capabilities of microbes based on genomic data alone. PROTEAN generates constituent metabolic production networks with maximum parsimony and probability to predict the production of a given metabolite with a defined set of input metabolites. PROTEAN is a combination of well-validated methods, including Parsimonious Enzyme Usage Flux Balance Analysis (pFBA) (37), likelihood-based gap filling (44), fastGapFill (45), and CarveMe (46). The algorithm uses the ModelSEED biochemical reaction database, a large set of known metabolic reactions, for constituent network generation (47). First, reaction likelihoods are calculated for each reaction in the ModelSEED database using Probannopy (48) (Figure 2). Reaction likelihoods correspond to the probability that a given reaction is catalyzed by an enzyme that is encoded for by the genome. We modified pFBA to utilize reaction likelihoods for weighted minimization of flux through each reaction, while still maintaining near-optimal flux through the objective function. Standard pFBA assumes that metabolism is optimized to minimize enzymatic turnover and thus the method is driven by a minimization of the total flux through the metabolic network (37). Weighted pFBA allows for the reconstruction of constituent anabolic networks while accounting for maximum genomic probability and resource parsimony (see Methods). The constituent anabolic networks output by PROTEAN consist of flux-carrying reactions required for the production of a certain metabolite with preferential flux through reactions that have higher reaction likelihoods. A constituent network represents a theoretically optimal biosynthetic network while accounting for the greatest genomic evidence for production of a given metabolite in a set media condition (Table S4). We represent the information from each constituent network using a single summary metric referred to as the Production Likelihood by calculating the average of all likelihoods of reactions that carry flux. The average of all reaction likelihoods in a metabolic pathway has been previously shown to be a valuable metric for making comparisons between networks (44).

**Figure 2:**
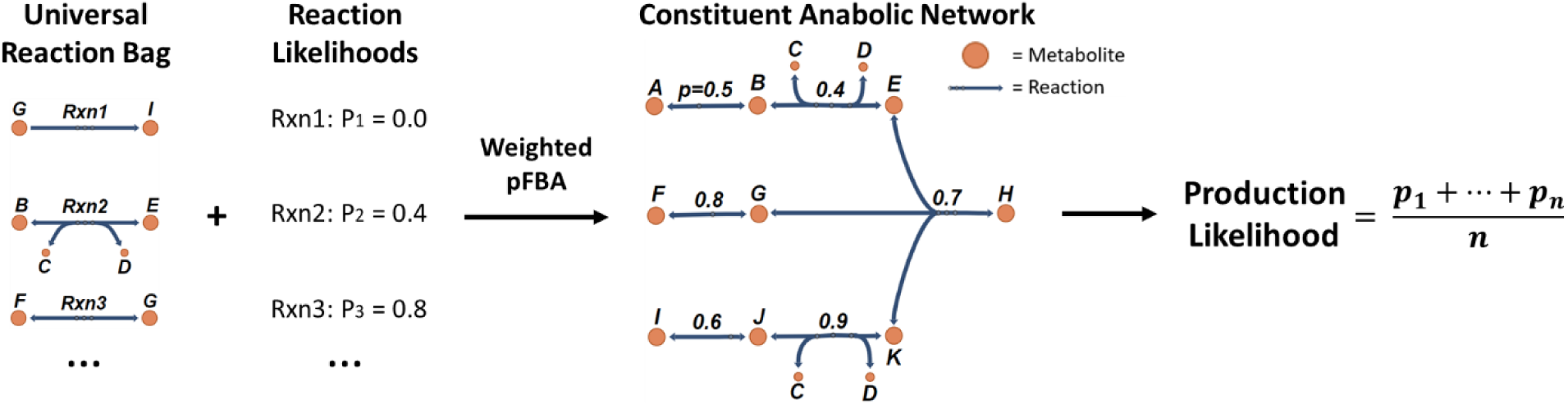
PROTEAN is an approach for quantifying the likelihood that a given metabolic network, derived exclusively from genomic evidence, is capable of synthesizing a particular metabolite. A modified version of Parsimonious Enzyme Usage FBA (weighted pFBA) was performed on a standardized set of reactions to generate constituent anabolic networks for each genome. Reaction likelihoods were used to weight the minimization of flux through each reaction in the network. Therefore, reactions with a greater likelihood were more likely to be included in the resulting constituent anabolic network. Each constituent network has a set of input metabolites representing the media condition (Table S4) and a demand reaction for a certain metabolic product. The resulting constituent network is the set of reactions that requires flux to produce the metabolic product in the given media condition. The production likelihood metric is an average of all the reaction likelihoods associated with the reactions included in the constituent network. This metric is used as a summary statistic that allows for the comparison of constituent networks across different metabolic products and strains, where a higher production likelihood corresponds with greater genetic evidence for that particular constituent anabolic network.

### The Scaled Production Likelihood metric facilitates comparison of anabolic capabilities between species and strains

Predicted constituent anabolic networks were generated for a set of 50 biologically-relevant metabolic products for each of the 144 *Lactobacillus* strains. The 50 metabolites were selected based on known *Lactobacillus* biology (see Methods). For each metabolic product, we generated a constituent anabolic network (Table S3) across all strains. For each genome we scaled the Production Likelihoods metric by calculating the corresponding z-score. The standard deviation for the z-score calculation was across all metabolic products for each strain. This metric allows for a relative comparison of production capabilities across strains that does not rely on well-curated metabolic network reconstructions. The resulting Scaled Production Likelihood (SPL) is a metric indicating likelihood that a genome encodes for the cellular machinery required to produce a metabolite, given a specific media condition, relative to all of the other SPLs for the metabolic products per strain. For visualization, these data were grouped by species and summarized using the median of the SPLs across all of the strains within each species (Figure 3).

**Figure 3:**
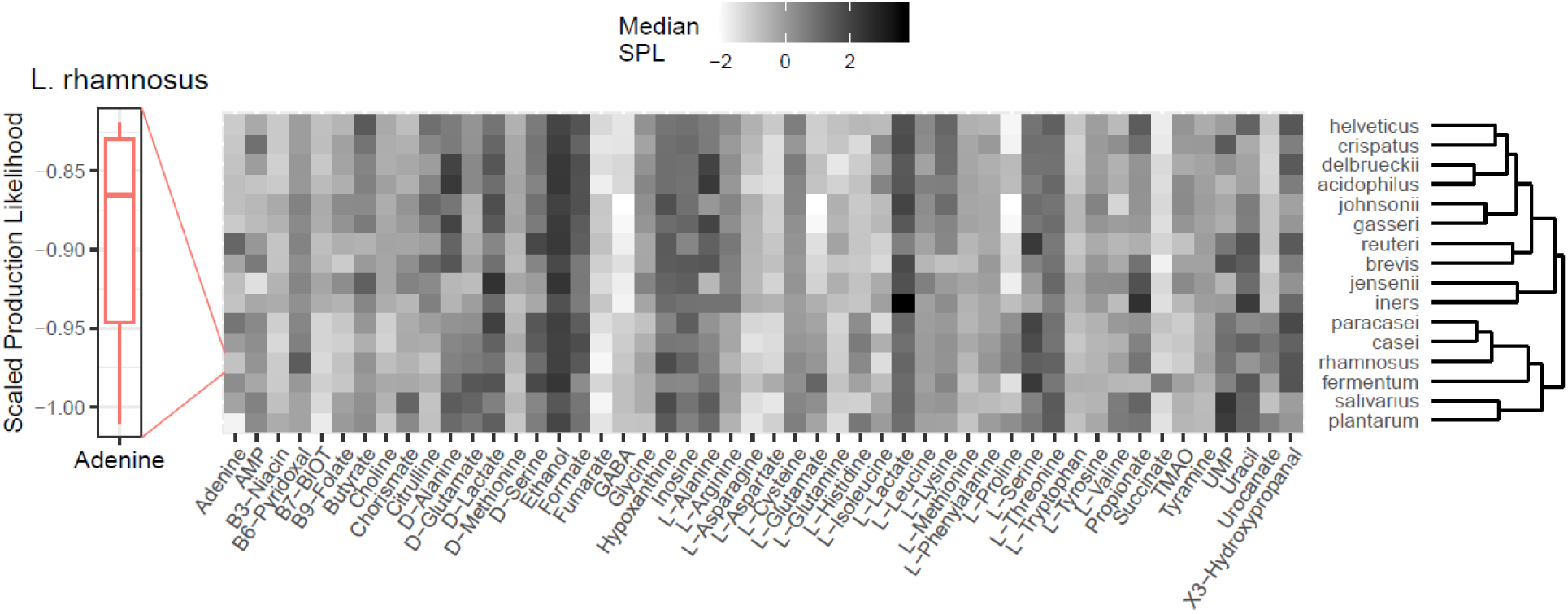
Predicted metabolic production capabilities with the Scaled Production Likelihood (SPL) metric align poorly with phylogeny. There is a single production likelihood for each genome associated with each metabolite. A median SPL can be calculated for a species that allows for more general comparisons across species, illustrated here by the distribution for one species (*L. rhamnosus*) and one metabolite (adenine). There are 50 metabolites used as features to allow for the comparison of predicted production capabilities across the lactobacilli analyzed.

The strains were grouped by species and clustered based on median SPLs. We found that across the 16 species, D- and L-lactate both have high median SPLs, as we would expect with lactobacilli. Additionally, fumarate and GABA have particularly low SPLs across all species. We were able to find several publications indicating GABA can be produced by select lactobacilli in specific environments (49,50). However, we were unable to find publications discussing the production of fumarate by lactobacilli. Additionally, we found that the dendrogram from clustering based on predicted metabolic production capabilities does not qualitatively align well with published phylogenetic trees generated using the 16S rRNA gene (34). The misalignment to established phylogenetic trees indicates that phylogeny is a poor indicator of metabolic production capabilities. It is likely that evolution of metabolic production capabilities is driven independently from classical genes used for phylogenetic comparisons, such as the 16S rRNA gene. Therefore, we need more precise computational tools to better understand the phenotypic differences between microbial species when interrogating metabolism. Perhaps phylogenetic analysis would be augmented with the consideration of metabolic genes in addition to the 16S rRNA gene.

### Intestinal and oral *Lactobacillus* strains have different metabolic capabilities compared to vaginal strains

We performed principle coordinate analysis (PCoA) on the SPLs for each species and determined that the *Lactobacillus* strains cluster significantly by both species (Figure 4A) and isolation site (Figure 4B) (PERMANOVA; *P* < 0.001). The vaginal isolates differ from both the oral and gut cluster (Figure 4B). Substantial overlap was found between oral and gut isolates, specifically within *L. gasseri, L. rhamnosus*, and *L. salivarius*, likely due to the consistent transmission of orally colonized microbes to the intestines (15). It has been hypothesized that many of the lactobacilli isolated from the gut are actually transient strains that are colonized in the oral cavity (51). Our data supports this hypothesis by showing that oral isolates are metabolically similar to a portion of the intestinal isolates. However, there are lactobacilli, such as *L. reuteri*, which likely colonize the human intestines (52). Five of the 16 species in this study are only represented by strains isolated from the intestines; although this result is influenced by sampling bias in the PATRIC Database, it provides support that our data contains species that are only found in the intestines. The vaginal isolates cluster separately from the intestinal/oral isolates along the primary coordinate that accounts for 78% of the variation in these data. The vaginal microbiota is frequently dominated by several *Lactobacillus* species, such as *L. iners, L. crispatus*, and *L. jensenii* (53–55). This separation of vaginal isolates from intestinal/oral isolates indicates that these two main clusters have differences in their metabolic production capabilities. This result is to be expected because the intestinal/oral nutrient environment is drastically different from the vaginal environment and the dominant species appear to have metabolic capabilities that reflect this difference.

**Figure 4:**
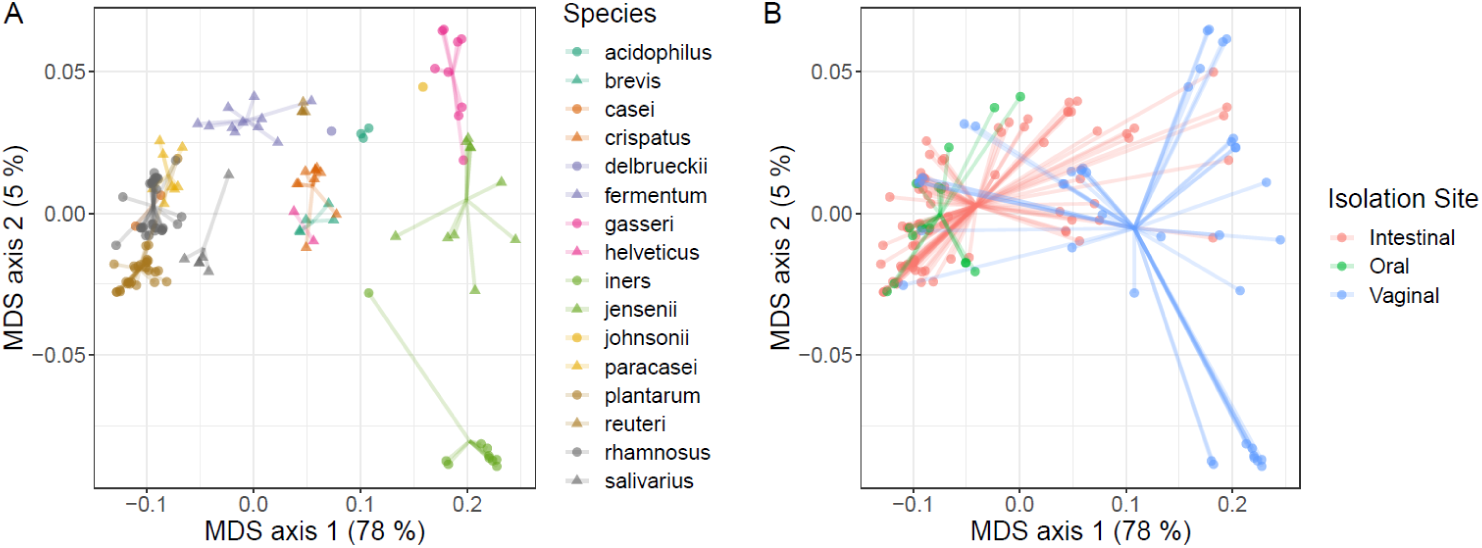
The Scaled Production Likelihood metric distinguishes metabolic functionality among species. (A) We found that *Lactobacillus* strains cluster significantly by species (PERMANOVA; P < 0.001). (B) Additionally, they cluster significantly by isolation site (PERMANOVA; P < 0.001). Both plots are PCoA using the Bray-Curtis distance metric of the SPLs for each isolate. Points in both panels are identical, but displayed with different color schemes.

In addition to distinguishing isolates by body site, the SPL metric is capable of defining collections of functional components that drive differences between groups. Using standard genomic analyses, differences between groups are typically defined by the differential gene content. Genes are intrinsically part of a larger network of metabolism where absence of specific functionality related to a gene may be compensated for within the system. Since our approach is based on Production Likelihoods of specific metabolites, it functions within a more complex metabolic framework compared to the analysis of genomic data without the network context. Using machine learning, we were able to identify the set of metabolites for which each group of strains is more likely to encode the cellular machinery required for production. We conducted a machine learning feature selection to determine the metabolites that are most likely to be produced by each group of strains, intestinal/oral strains and vaginal strains. We grouped the intestinal and oral strains together due to their inherent similarity (Figure 4B) and the observed transmission of oral strains to the intestines (15,51). We generated two separate area under the curve random forest (AUCRF) models to determine the metabolites that were more likely to be produced by each of the groups. Two models were necessary to enrich for the most discriminatory metabolites that were more likely to be produced in each of the groups, rather than simply identifying the metabolites that best classify the samples based on isolation site regardless of being more or less likely to be produced (See methods). The first model was generated to select the metabolites that are most likely to be produced by the intestinal and oral isolates compared to the vaginal isolates, while maximally discriminating the groups. The eight metabolites selected accurately classify greater than 90% of isolates to the correct group (Figure 5A). The second model was generated to select the metabolites that are most likely to be produced by the vaginal isolates compared to the intestinal and oral isolates, while maximally discriminating the groups. The seven metabolites selected accurately classify greater than 90% of the isolates to the correct group (Figure 5B).

**Figure 5:**
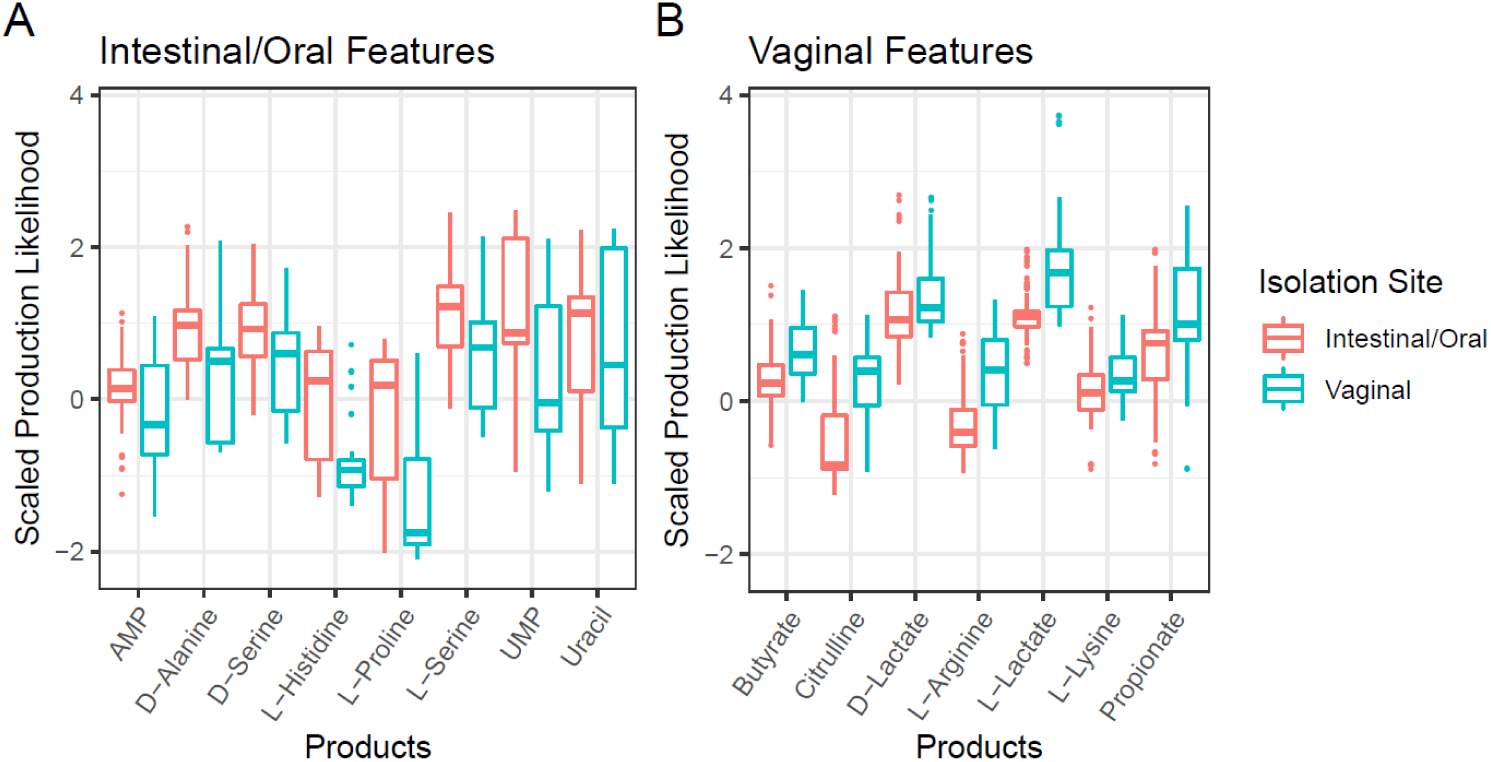
Machine learning of the SPL scores identifies metabolites that discriminate *Lactobacillus* strains. Machine learning feature selection identified the metabolites that are both most likely to be produced by each group and capable of classifying the strains into two groups, intestinal/oral and vaginal, with greater than 90% accuracy. (A) There are eight metabolites that are more likely to be produced by the intestinal/oral isolates compared to the vaginal isolates. (B) There are seven metabolites that are more likely to be produced by the vaginal isolates compared to intestinal/oral isolates. Both models are more than 90% accurate in predicting the membership to which the given isolate belongs using the SPLs of the metabolites listed.

Using SPLs as an input for AUCRF feature selection, we identified the metabolites that are most likely to be produced by the strains associated with the two isolate groups, intestinal/oral and vaginal. The selected metabolite products may contribute to how the strains interact with the mucosal tissues in each site. We hypothesize that these metabolites are related to key phenotypic differences between the two isolate groups. Four of the selected metabolites that are likely produced by intestinal/oral strains, D-alanine, D/L-serine, and L-proline (Figure 5A), have all been previously identified to have an impact on the human intestinal epithelium (23,24,56–58). Additionally, four of the selected metabolites that are likely produced by vaginal strains, L-arginine, citrulline, and D/L-lactate (Figure 5B), have been previously identified to have an impact on the human vaginal microbiome (59–62). The metabolites for which we have not found existing experimental evidence for are likely worth focusing on in future experimental studies.

For intestine-associated lactobacilli in this study, there is a connection between intestinal immune system regulation and D-alanine rich lipotechoic acid, a glycolipid expressed by some lactobacilli, such as *L. plantarum* (23,24). D-alanine rich lipotechoic acid, produced by lactobacilli, has been shown to down-regulate local colonic inflammation in a murine colitis model (23,24). With PROTEAN we identified that intestinal lactobacilli were more likely to produce D-alanine (Figure 5A). It is possible that a positive interaction with the intestinal host immune system would result in an evolutionary advantage by reducing local immune response. Additionally, serine rich serine-threonine peptides have been shown to have a similar regulatory effect on intestinal dendritic cells (56,57). These peptides expressed by *L. plantarum* are resistant to intestinal proteolysis and appear to be present in the colon of most healthy individuals (56,57). Similar to D-alanine, the production of D/L-serine would require a robust biosynthesis pathway present in those strains.

A final gut-related connection involves the biosynthesis of L-proline (Figure 5A). One of the primary stress responses in *L. acidophilus* to high osmotic pressure results in the accumulation of L-proline in the cell; there is little evidence that this response is a result of L-proline transport into the cell (58). These *Lactobacillus* strains are exposed to a large range of stressors in the gut, including suboptimal osmotic pressures. There is strong evidence that L-proline is used by *L. acidophilus* to tolerate suboptimal osmotic pressures and there is a lack of evidence for L-proline transporters. As such, the biosynthesis of L-proline may be advantageous for growth in the gut.

For the enriched metabolic products in vaginal isolates (Figure 5B), there is evidence for an arginine/ornithine antiporter and arginine deiminase in *L. fermentum* (59). These enzymes are part of the arginine deiminase pathway through which there is the production of citrulline which is exported from the cell and contributes to acid tolerance (59). It has also been demonstrated that treatment with probiotics containing arginine deiminase-positive lactobacilli can improve clinical symptoms of vaginosis in parallel with significant declines in polyamine (i.e. arginine, ornithine, and citrulline) levels in the vagina (60,61). The vaginal isolates in this study show enrichment for the cellular machinery required for the production of both citrulline and L-arginine (Figure 5B). The importance of lactate for the adequate maintenance of vaginal health in many individuals is known. The current hypothesis revolves around colonization resistance where vaginal lactobacilli establish an acidic environment by producing lactate (62). The acidic environment is generally inhospitable to invading pathogens as well as other microbes that are otherwise capable of residing in the vaginal environment (62). It has been shown that higher levels of D-lactate over L-lactate present in the vagina, produced by lactobacilli, further decrease the chance of infections in female patients (62). However, both isoforms of lactate remain important in maintaining vaginal health.

## Conclusions

Microbial biosynthesis of metabolites has a broad range of applications, from bio-manufacturing to microbiome research (63). There is a wealth of well-curated and accessible knowledge stored in biochemical reaction databases such as ModelSEED (64). Genome-Scale Metabolic Network Reconstructions access this fundamental knowledge while accounting for systems-level interactions. This study represents one such application of GENREs that is a step toward predicting the metabolic production capabilities of understudied organisms. Experimental validation of the production capabilities predicted with PROTEAN will allow for conclusions to be made beyond the statement that a microbe is genetically likely to be able to produce a metabolite. Utilizing PROTEAN data, we found that human-associated lactobacilli strains cluster significantly by species and isolation site. Additionally, many of the metabolic products that drive the clustering of strains by the isolation sites have known physiological function and importance in the respective isolation sites.

Future applications of PROTEAN could include optimal strain selection for bio-manufacturing of a certain compound, generating predicted metabolomics data for an organism to generate a prioritized list of conditions that would be most worthwhile to validate experimentally, and predicting the metabolites that are most likely to be produced in a microbiota. Microbes can have a wide range of physiological impacts on human health; these impacts are, in part, a result of the metabolites that are or are not produced by members of a microbiota. One of the core limitations of this study includes the lack of reaction likelihoods for some reactions in the universal reaction bag we used from ModelSEED. The number of reactions we could generate likelihoods for was limited by the Probannopy reaction template. However, this template can be expanded to continue to improve the utility of PROTEAN. With the inclusion of validation data, additional analyses will be possible, such as determining metabolic production pathways lacking proper annotation. By determining the reactions that are most likely required for biosynthesis of a known product, it would be possible to generate additional hypotheses for enzyme annotation experiments. PROTEAN is an algorithm with potential for a wide range of applications in the study and use of microbial metabolic networks.

## Methods

### Constituent Anabolic Network Generation (PROTEAN)

Probabilistic pFBA-based constituent anabolic network generation was accomplished using three Python packages, Cobrapy (65), Mackinac (66), and Probannopy (48). The complete ModelSEED universal reaction bag was downloaded from the github repository and filtered based on the annotation quality score, including all reactions with an ‘OK’ quality status or better (64). For each reaction in the ModelSEED universal reaction bag, we used Probannopy to generate a reaction likelihood based on the FASTA file for each genome obtained from the PATRIC database (4). The Cobrapy implementation of Parsimonious Enzyme Usage Flux Balance Analysis (pFBA) was altered to allow for each reaction’s linear constraint to be set individually based on the reaction likelihood. The linear constraint for each reaction was set to one minus the reaction likelihood (a value between 0 and 1). There were reactions included in the universal reaction bag that were lacking from the Probannopy template model, therefore resulting in several gene-associated reactions lacking reaction likelihood scores. The reactions without likelihoods were left at a full minimization penalty (linear constraint value of 1). We chose to penalize the reactions without likelihoods to bias our results towards the construction of networks for which all reactions had evidence of presence. The linear constraints applied to each reaction based on likelihood acted as a weighting (inclusion penalty) for the minimization step in pFBA, resulting in the reactions with greater likelihood having a lower penalty for carrying flux; therefore, the reactions had a higher likelihood of being included in the constituent anabolic networks.

Using PROTEAN, we generated constituent anabolic networks by setting a certain input media condition (Table S4) and constraining flux through the single metabolite objective function (Table S3). We ran our likelihood-weighted pFBA flux minimization across the entire universal reaction bag and isolated the reactions that carried flux to get the desired product. The resulting networks consist of the direct reactions that would be part of a production pathway as might be shown in a typical biosynthesis pathway figure, while also accounting for all of the secondary and energy metabolites that are required for the production of the metabolite in consideration. Additionally, this algorithm is optimizing for three core characteristics in the constituent networks: 1) minimum flux through the network (loosely, the minimum number of reactions), 2) maximum average reaction likelihood across the constituent network, and 3) output flux within 90% of the optimal yield of the metabolic product. We chose to allow flux through any reaction in the universal reaction bag during the generation of the constituent anabolic production pathways rather than simply pulling from a GENRE that was first gapfilled to allow production of biomass. Using the universal reaction bag instead of a gapfilled model was important because the biomass function is difficult to define for understudied organisms and unnecessary for our applications.

### Scaled Production Likelihood Metric

We represent the information from each constituent network using a single summary metric for ease of comparison, simply named the Production Likelihood. This metric is the average of the reaction likelihoods included in the constituent network. The average reaction likelihood for a metabolic pathway has been previously used for making comparisons between networks (44). The Production Likelihoods for all 50 metabolites are scaled for each given genome by calculating the z-score to create the Scaled Production Likelihoods used for the majority of the analysis in this study. The z-score is calculated for each individual strain using the median and standard deviation for the production likelihoods across the 50 metabolic products. The Scaled Production Likelihood allows for a ranked comparison of metabolic products across the genome set and corrects for annotation bias by essentially comparing the ranked z-score for each metabolic product.

### Supporting data for pathway generation

The simulated media formulation was based on *in vitro* minimal media growth conditions for *L. plantarum* (Table S4) (67–69). The techniques used in this study do not assume that all species are capable of growth in the given media condition, therefore this media condition simply provides a standard reference for comparison. The product list was developed by identifying metabolites that have been shown to be produced by lactobacilli during *in vitro* growth experiments, in addition to other metabolites that have been shown to be related to human physiology (70–74).

### Machine learning feature selection

Discriminating intestinal/oral and vaginal features were selected using area under the ROC curve random forest (AUCRF) using default parameters (75) (see Code). We generated two separate AUCRF models to determine the metabolites that were more likely to be produced by each of the groups, intestinal/oral and vaginal. Two models allowed us to enrich for likely products rather than simply selecting for the metabolites that provide the greatest discrimination between the groups but which may have poor likelihood scores. We conducted the enrichment for likely metabolic products for each model by reducing the feature set down to only metabolites that were more likely to be produced by the group of interest. Likely metabolic products were determined by comparing the median SPLs of each metabolite between the groups. Additionally, the feature sets were reduced to include only metabolites with a median value greater than zero for the group of interest. An AUCRF model was then generated to select the features that provided the greatest discrimination between the two groups.

### Statistical modeling and figure generation

The principle coordinate analysis (PCoA) ordinations were created using the R vegan package (76), implemented with the Bray-Curtis dissimilarity metric. Statistical significance for comparing the PCoA clusters was determined using a PERMANOVA (R Adonis test). A variety of R packages were used for all figure generation (77–81).

### Genome Quality and PATRIC Cross Genus Protein Family Data

Genomes used in the study were filtered for quality before being included in the analysis. Strains with greater than 0.2% unknown nucleotide calls in the genome were eliminated. Low quality genome assemblies with greater than 300 contigs were removed. Non-human associated *Lactobacillus* strains from the PATRIC database were used to determine the GC content range for each species (82,83), and significant outliers (plus or minus two percent) were removed to control for sequencing bias (84,85). Only isolates from the three human-associated sites (oral, intestinal, and vaginal) were included in the final dataset.

The inclusion of metabolic PATRIC cross genus protein families was conducted by filtering the PGfams for each genome based on the existence of an associated known reaction and Probannopy likelihood greater than 0. Pan and core metabolic PGfam sets were evaluated after the addition of all genomic features from each genome. The pan set of metabolic PGfams was defined as the total number of unique PGfams included in the data set after the above filtering steps. The core set of metabolic PGfams are those that existed within each genome included in this study.

## Data and code availability

Genome FASTA files and metadata were downloaded from the PATRIC Database (4). Python and R code is available at: Github.com/Tjmoutinho/Lactobacillus

